# Curved Axially Scanned Light-Sheet Microscopy

**DOI:** 10.64898/2025.12.03.691599

**Authors:** Steven J. Sheppard, Tatsuya C. Murakami, Douglas P. Shepherd

## Abstract

Light-sheet fluorescence microscopy enables high-throughput multidimensional imaging. However, attempts to further improve throughput by incorporating commercially available or custom high space-bandwidth product objectives have been limited by objective field curvature, which violates the co-planar overlap required for light-sheet imaging and ultimately reduces the usable field of view. Inspired by machine-vision strategies that curve the image plane to match the field curvature, we introduce curved axially scanned light-sheet microscopy, which adapts the light-sheet excitation to the detection objective’s field curvature via synchronized control of the remote refocus scan and motorized mirror. Using our technique, we increase the usable field of view along the light-sheet propagation axis from ∼2.5 mm to ∼6.3 mm for a commercial high-SBP detection objective with significant field curvature, while maintaining sub-cellular resolution.

## 1. Introduction

Selective plane illumination microscopy (SPIM), otherwise known as light-sheet fluorescence microscopy (LSFM), has become an essential tool in biological imaging, enabling high-resolution volumetric imaging with optical sectioning and reduced photo-toxicity compared to conventional wide-field or confocal modalities [1, 2]. By using orthogonal, co-planar detection and excitation volumes confined to a relatively thin region, LSFM minimizes photo-bleaching and enables rapid acquisitions of large 3D volumes [3]. Using LSFM to increase throughput, in particular the total volume imaged per unit time, enables new insights into fast biological dynamics as well as rapid imaging of large cleared samples [4–8]. Despite innovations in microscope objectives, large samples often require tiling multiple areas because the usable field of view (FOV) is limited to the region where the detection objective maintains spatially invariant resolution and the co-planar requirement is satisfied. Ultimately, improving the achievable size of one LSFM FOV is tied to the ability to effectively increase the area of co-planarity without loss of resolution or optical sectioning.

A useful theoretical metric of imaging system throughput is the space-bandwidth product (SBP), or effective information capacity,

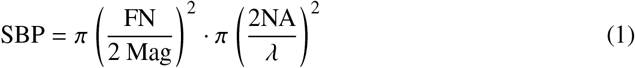

where the imaging objective’s field number (FN) and magnification (Mag) at the design tube lens focal length determine the theoretical field of view (FOV). The spatial frequency support of the objective’s numerical aperture (NA) at wavelength *λ* defines the objective’s maximum spatial resolution. Specifically, for incoherent imaging, the maximum spatial frequency is 2NA *λ* [9].

In reality, it is challenging to build imaging systems that fully realize their theoretical SBP. Both the usable FOV and resolution can be limited by the camera’s pixel pitch and number of pixels, which must simultaneously satisfy the Nyquist-Shannon sampling criterion and provide sufficient pixels to capture all the information within the objective. Even with an appropriate camera, the area of spatially invariant resolution (governed by optical aberrations) also limits the usable FOV and achievable SBP. Moving from general optical systems to LSFM, the requirement for a co-planar excitation and detection volumes further constrains the usable FOV. To this end, LSFM method development has expanded the SBP by engineering sophisticated excitation and detection strategies. These include custom optics, strategies to maintain uniform optical sectioning across the FOV, engineering the shape of the light-sheet, and many others. Focusing on large sample imaging, existing efforts have combined axially scanned light-sheet microscopy (ASLM) with high-SBP objectives [10–14]. ASLM increases the SBP of the LSFM approach by synchronizing the sweep of the light-sheet propagation axis focus to the rolling shutter of a scientific CMOS (sCMOS) camera, increasing the area of invariant spatial resolution in all three dimensions [10, 15]. Because ASLM instruments often have a small excitation volume, this imposes stricter co-planarity requirements, and both heterogeneous refractive index (RI) and detection objective field curvature can limit the achievable SBP [16, 17]. RI heterogeneity is often addressed by reducing the excitation NA at the cost of optical sectioning, optimizing sample preparation to homogenize RI, using custom sample chamber geometries, or dynamically optimizing the remote focusing parameters [16–21]. However, field curvature limits the ASLM FOV to the area of co-planarity between the illumination and detection volumes or requires engineering bespoke solutions optimized for individual LSFM objective pairs [17]. In the machine vision community, a common strategy to account for field curvature is to use a curved image sensor [22] or build image-sensor arrays out of many individual cameras [23, 24]. The lack of a programmable shutter immediately rules out curved detector strategies for LSFM platforms that use the shutter as a virtual confocal slit, motivating alternative approaches to account for field curvature.

Some projects, such as the mesoSPIM initiative, have tested and selected the “best” commercial objectives with minimal field curvature [12]. Other approaches, such as the ExA-SPIM and the Mesolens, designed custom high-SBP objectives to minimize field curvature while minimizing RI heterogeneity via sample expansion [14, 25]. The Mesolens approach uses a global shutter pixel-shifting camera to increase pixel sampling, at the cost of imaging speed and ASLM operation [25]. curvedLSFM designed a custom objective and a custom illumination path to produce excitation volumes that match the field curvature of the detection volume perpendicular to the light-sheet propagation direction [26]. Because field curvature is matched exactly only at the focus of a propagating light-sheet in curvedLSFM, the method requires a line-integration camera and mechanical translation of the sample to build an image. Such an approach yields excellent-quality data, yet, because measurements are not parallelized across the light-sheet, curvedLSFM is closer to a large-format line-scanning confocal microscope than to an LSFM. Table 1 provides a comparison of the theoretical SBP (detection objective only using eq. 1) and effective SBP (detection optics, usable FOV due to field curvature, and camera properties) for select high-SBP LSFM approaches. Fundamentally, all of these approaches share the same goal: maintaining a co-planar excitation and detection volume while increasing throughput.

**Table 1.** High SBP LSFM approaches. The theoretical objective (Obj.) SBP accounts for the FOV at the specified NA but does not reflect the diffraction-limited FOV. For the Olympus MVPLAPO series objectives (MesoSPIM v5, CurvedASLM), we report the diffraction-limited FOV and SBP, followed by the geometric values (range of NA support) in brackets. The diffraction-limited FOV for the Olympus MVPLAPO 2x was estimated from Ref. [27], and the MVPLAPO 1× value is assumed to be twice the optical axis distance. The effective (Eff.) SBP was computed using the experimental FOV diagonal and reported experimental resolution, which captures the performance of both the excitation and detection pathways, including camera sampling. For curvedLSFM, a line-scanning camera is used, and therefore the Eff. SBP is ill-defined and not reported. Key: *-ASLM; ^-LSFM; †-line scanning confocal microscopy; *δ*-effective lateral resolution limited by camera pixel pitch.

| Microscope | Obj. NA | Obj. FOV (mm) | Obj. SBP ($\cdot 10^6$) | Eff. single FOV (mm) | Eff. resolution ($xy \times z$) ($\mu\text{m}$) | Eff. SBP ($\cdot 10^6$) |
| --- | --- | --- | --- | --- | --- | --- |
| mesoSPIM v5* [11] | 0.25 | 4.5 [34] | 46 [2637] | 3.3 x 3.3 | 3.25 x 5.0<sup>δ</sup> | 2.5 |
| Benchtop mesoSPIM* [12] | 0.14 | 4.8 | 16.5 | 5 x 3.5 | 2.6 x 3.3 | 9.1 |
| Airy Mesolens<sup>$\wedge$</sup> [13] | 0.47 | 6 | 290 | 4.4 x 3 | 0.55 x 7.8 | 246 |
| ExaSPIM* [14] | 0.305 | 16.8 | 958 | 10.6 x 8.0 | 1.5 x 2.5<sup>δ</sup> | 193 |
| CurvedLSFM<sup>†</sup> [26] | 0.25 | 13 | 385 | 10.24 x 0.08 | 1.25 x 2.5<sup>δ</sup> | — |
| CurvedASLM* | 0.5 | 2.25 [17] | 46 [2637] | 6.29 x 4.8 | 2.8 x 2.5<sup>δ</sup> | 19.6 |

In this work, we present curvedASLM, a method to increase the FOV for any pair of LSFM objectives and immersion media by curving the light-sheet path along the curved high-contrast plane of the detection optics. curvedASLM is fundamentally different from the other approaches in Table 1 because we dynamically generate the correction based on a simple calibration. More specifically, we synchronize an electrotunable lens and a galvanometric mirror to perform ASLM while also translating the beam along the detection axis. By matching the light-sheet to field curvature, curvedASLM addresses the hardware and optical limitations that limit many high-SBP LSFM implementations. We demonstrate an SBP improvement from 9.1 to 19.6 for a prototype platform using a commercially available high-SBP detection objective with significant field curvature [27]. Using this platform, we imaged contrast targets in various refractive index media to calibrate media-specific field curvature and fluorescent microspheres to quantify the gain in usable FOV. We then demonstrated improved image quality by imaging the vascular network of a whole mouse brain with and without a curved light-sheet. Because curvedASLM increases the usable SBP by controlling the light-sheet rather than implementing custom detection optics or curved digital sensors, we anticipate that this approach can improve existing ASLM installations with minimal modification.

## 2. Materials and Methods

### 2.1. Experimental Setup

We modified an existing ASLM [21] to control the light-sheet position along the focus axis of the detection pathway, as illustrated in Fig. (2). The main change was the introduction of two additional lenses to conjugate a large galvanometric mirror to the EO_bfp_. The control computer was a Lenovo P360 running Ubuntu 22.04.5 LTS (Intel Core i7-12700, Nvidia RTX 3060 GPU, 64 GB RAM) with a digital acquisition card (National Instruments PCIe-6343 with two SCB-68A connectors). We used a modified version of the open-source navigate software to synchronize the hardware and acquire data [28, 29]. We provide a detailed description of the excitation and detection pathways below.

**Fig. 1.**
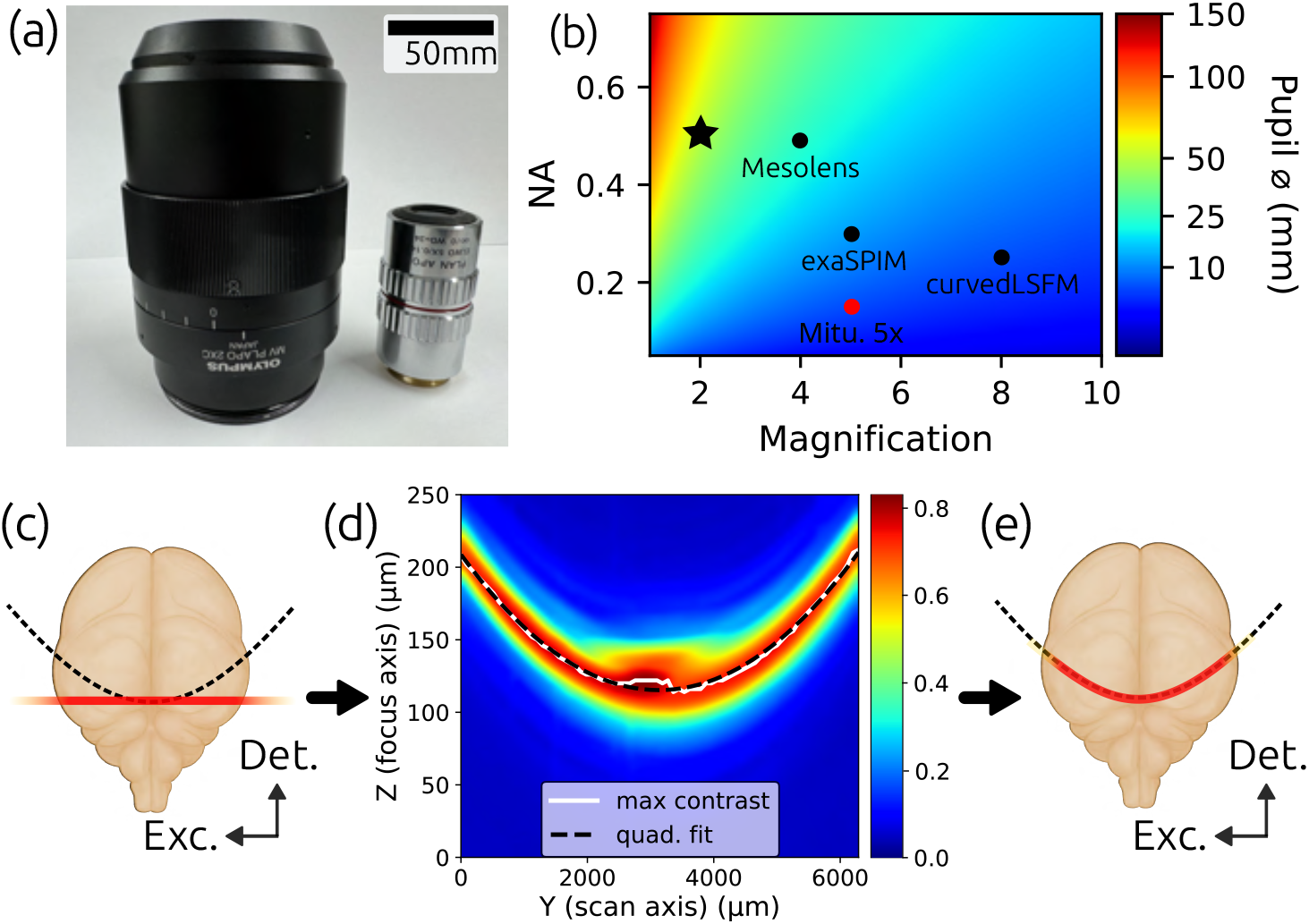
CurvedASLMoverview. (a) SizecomparisonbetweentheOlympusMVPLAPO 2×(NA0.5)andMitutoyo5×(NA0.14)longworkingdistanceobjectives. (b)Heatmap of the Pupil diameter for a given lens NA and magnification, assuming a 100mm tube lens. The objective used in our work, Olympus MVPLAPO 2X (star) and other large sample light-sheets are shown, along with the Mitutoyo 0.14 NA objective for reference. (c) Discrepancy between excitation (exc.) and detection (det.) pathways in conventional LSFM/ASLM. (d) Characterizing the detection pathway’s curved plane of maximum contrast. (e) Maintaining coplanar alignment of the excitation and detection pathways with curvedASLM.

**Fig. 2.**
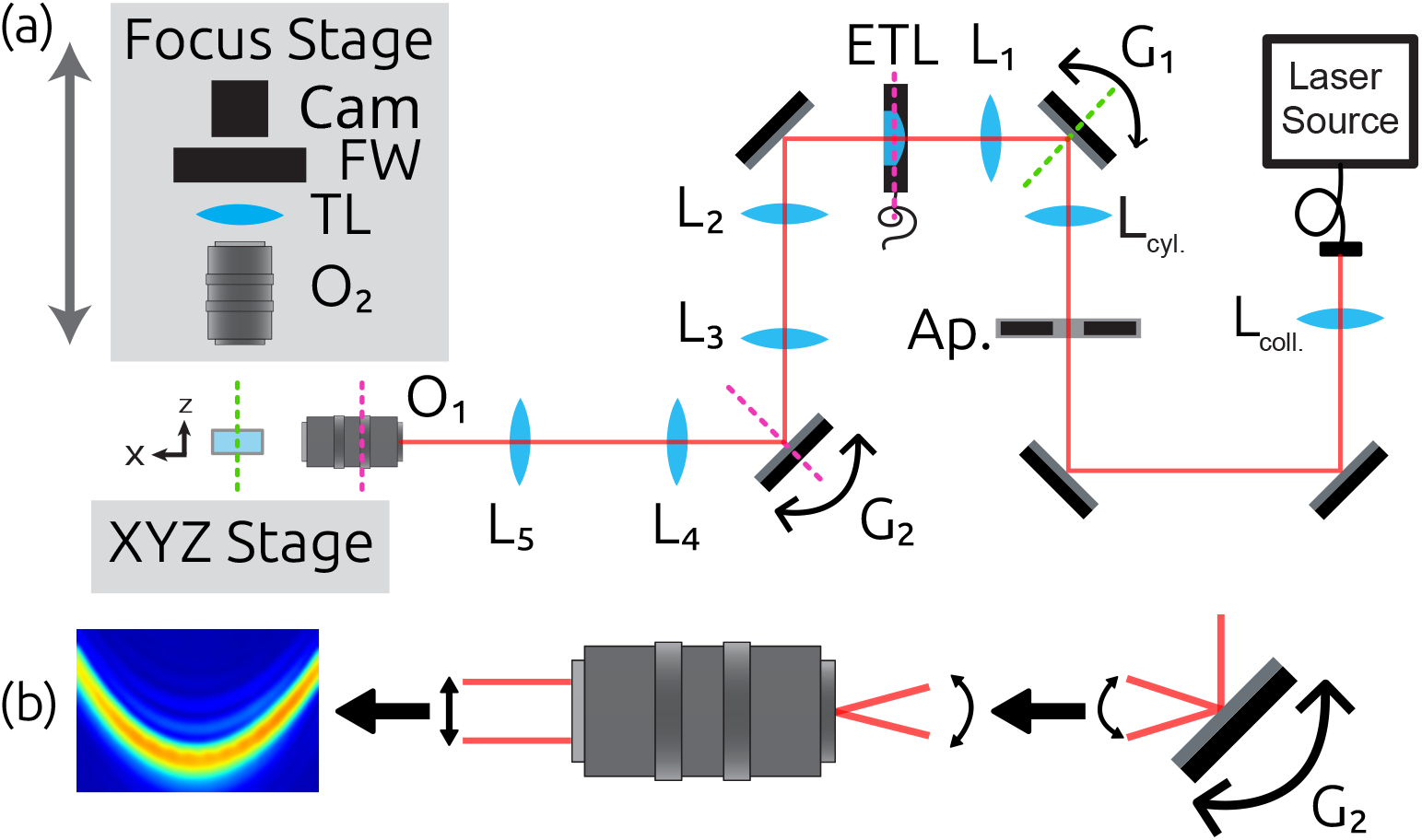
CurvedASLM prototype platform. (a) Experimental optical pathway. L_i_ are achromat doublets, Ap. is the adjustable aperture used to modulate the NA. L_cyl_ is the cylindrical lens, and TL is the tube lens. The magenta & green dotted lines represent conjugate planes to the BFP_EO_ and FFP_EO_ respectively. (b) Follow the maximum contrast by modulating G_2_’s angle to translate the light-sheet in the FFP_EO_ along the z-axis.

The excitation pathway consists of a laser source (Oxxius L4Cc) with four wavelengths (473 nm, 532 nm, 561 nm, 638 nm) combined into a single fiber output. The output was collimated with a 200 mm achromatic lens (Thorlabs AC508-200-ML) and the effective NA was tuned using an adjustable rectangular slit (Ealing 74-1137-000) conjugate to the EO_bfp_. The light-sheet profile was generated by a 50 mm cylindrical lens (Thorlabs ACY254-50) which was focused on a 5 mm galvanometric mirror (Thorlabs GVS201) to enable pivoting the light-sheet for shadow reduction [30]. The beam is collected by a 50 mm lens (Thorlabs AC254-50-ML) and focused on an electrotunable lens (ETL)(Optotune EL-16-40-TC-VIS-20D-1-C and Gardasoft TR-CL180 controller) used for remote focusing. A two lens relay (Thorlabs AC508-180-ML, AC508-100-ML) conjugated the ETL onto the focus shifting galvanometric mirror (Thorlabs QS30X-AG). A second two lens relay (Thorlabs AC508-100-ML, AC508-150-ML) conjugated the focus shifting mirror to the back focal plane (EO_bfp_) of our 0.14 NA and 5× magnification excitation objective (Mitutoyo 378-802-6). Using the rectangular slit, the light-sheet height was tunable up to ∼4800 µm.

The detection pathway consists of a large life sciences detection objective (Olympus MVPLAPO 2X) and “tube lens” (Olympus MVPLAPO 1X), yielding a 0.5 NA and 2× magnification. The detection objective has a nominal FOV of 17 mm, a working distance of 2 cm, and a back focal length of ∼5 mm. To fully capture the ∼50 mm pupil of the 2× objective, we machined a custom mount to position the objective and tube lens with a ∼10 mm separation. We used a sCMOS camera with 4.25 µm pixels and programmable rolling shutter (Teledyne Photometrics Iris15). With an overall 2× magnification, we had an effective pixel size of 2.125 µm and camera limited FOV width of 6290 µm. The effective pixel size does not satisfy Nyquist-Shannon sampling requirements, resulting in pixel-limited resolution of 2.125 µm. The fluorescence excitation was isolated with a 30 mm diameter filter wheel positioned between the tube lens and camera (Finger Lakes Instrumentation HS0433417, Filters: Semrock FF01-536/40-32-D, FF01-624/40-32-D). The entire detection pathway was mounted on a motorized breadboard (Thorlabs KST101, MB1012), which was adjusted to maintain focus. Samples were mounted on a *XYZ* motorized stage for z-stacking and multi-position imaging acquisitions (3× Applied Scientific Instrumentation LS-50 stages configured for *XYZ* motion and Applied Scientific Instrumentation MS-2000 controller).

### 2.2. Characterizing the detection pathway contrast

To characterize the detection pathway’s focus contrast, we adapted the method in ref [12]. We mounted a Ronchi ruler with 80 lp/mm (Thorlabs R1L3S15N) to the *XYZ* stage, back-illuminated with a diffuse white light source, and acquired z-stacks in multiple imaging media. The contrast C was calculated as follows.

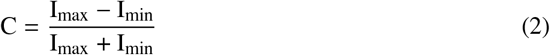

### 2.3. Mapping the light-sheet to the detection field curvature

We synchronized the light-sheet position with both the camera’s rolling shutter and the detection pathway’s plane of maximum contrast across the entire rolling shutter axis of the camera chip. To calibrate the lateral shift (relative to the light-sheet propagation direction) and applied voltage to the galvanometric mirror, Δf /ΔV, we measured the light-sheet offset using a fluorescing cube over a voltage range of ±1 V. We further validated this measurement in a fluorescent microsphere sample by adjusting the light-sheet position until the lateral PSF at the edges of the camera FOV was restored to near-diffraction-limited. Contrary to the Ronchi ruler measurements, we found that the required voltage from one edge of the FOV to the other differed by 10%, requiring a third-order polynomial rather than a quadratic waveform. We suspect that the rotational degree of freedom in our sample mounting caused a symmetric contrast map with the Ronchi ruler. Our current approach is to manually tune the voltage for the FOV edges, along with the remote focus sweep parameters, for each sample.

### 2.4. Data processing

The acquired data follows a curved path, which results in multiple z-planes within a single image. Similar to ref. [26], the axial displacement throughout the image has the linear relationship Δz (x) = df /*dV*.1/V (x) . We developed a Python framework that applies a linear shift based on the applied waveform, fuses multi-tile acquisitions into a ome-zarr v0.4 datastore [31], and optionally performs computational de-striping [32, 33].

### 2.5. Sample Preparation

#### 2.5.1. Fluorescent microspheres in agar

We prepared 1.0 µm yellow-green (YG) fluorescent microspheres (Invitrogen F8823) embedded in 1 % low melting agar. The YG microsphere stock was diluted in water to 10^−4^, and sonicated for 20 min. We then added 1 mL of the diluted beads to 4 mL of the 1 % low melting agar (Sigma-Aldrich A4018-70G) at 60 ^°^C and mixed using pipette action. The liquefied agar was transferred to a 10 mm×20 mm cuvette (Starna Cells 3-Q-20) and allowed to cool before imaging.

#### 2.5.2. Mouse Brain

All procedures were approved by The Rockefeller University Institutional Animal Care and Use Committee (IACUC) and were in accordance with the National Institutes of Health guidelines. The 8-week-old mouse was sacrificed by an overdose of pentobarbital (>100 mg kg^−1^), then transcardially perfused with 15 mL of phosphate-buffered saline (PBS) and 20 mL of 4 % paraformaldehyde (PFA, Electron Microscopy Sciences) in PBS. The brain was dissected and post-fixed with 40 mL of 4 % PFA in PBS at 4 ^°^C overnight. The fixed tissue was transferred directly into an 80 % (v/v) methanol (MeOH) solution in water at 4 ^°^C and gently shaken for 2 hours for dehydration. Dehydration was repeated using 80 % (v/v) MeOH with shaking for 2 hours at 4 ^°^C. The sample was then transferred to 100 % MeOH with shaking for 2 hours at 4 ^°^C. After refreshing the MeOH, the sample was moved to room temperature (RT) and shaken overnight. The MeOH was refreshed again, and the sample was transferred to 37 ^°^C with shaking for 5 days.

The sample was rehydrated using a decreasing gradient of MeOH. The concentrations and durations were 80 % for 30 min, 50 % for 30 min, and 30 % for 30 min. After rehydration, the iDISCO protocol was followed until clearing [34]. In this study, a 1/1000 dilution of Cy3-conjugated anti-αSMA (C6198-100UL, Sigma-Aldrich) was used. The sample was cleared with BABB (1:2 mixture of benzyl alcohol and benzyl benzoate) after MeOH dehydration.

The sample was imaged in a 10 mm ×20 mm cuvette (Starna Cell 3-Q-20) using a 3D printed mount [11, 12] and immersed in BABB.

## 3. Results

### 3.1. Field curvature characterization

We calculated the contrast in 40× 40 pixel regions of interest (ROIs) and used the 99th percentile for the maximum and minimum intensity values. A summary of our results is presented in Fig. 3. We characterized the detection optics’ contrast in air (n =1.0), and with a large outer refractive index matching cuvette filled with water (n =1.33) or microscopy oil (n =1.515).

**Fig. 3.**
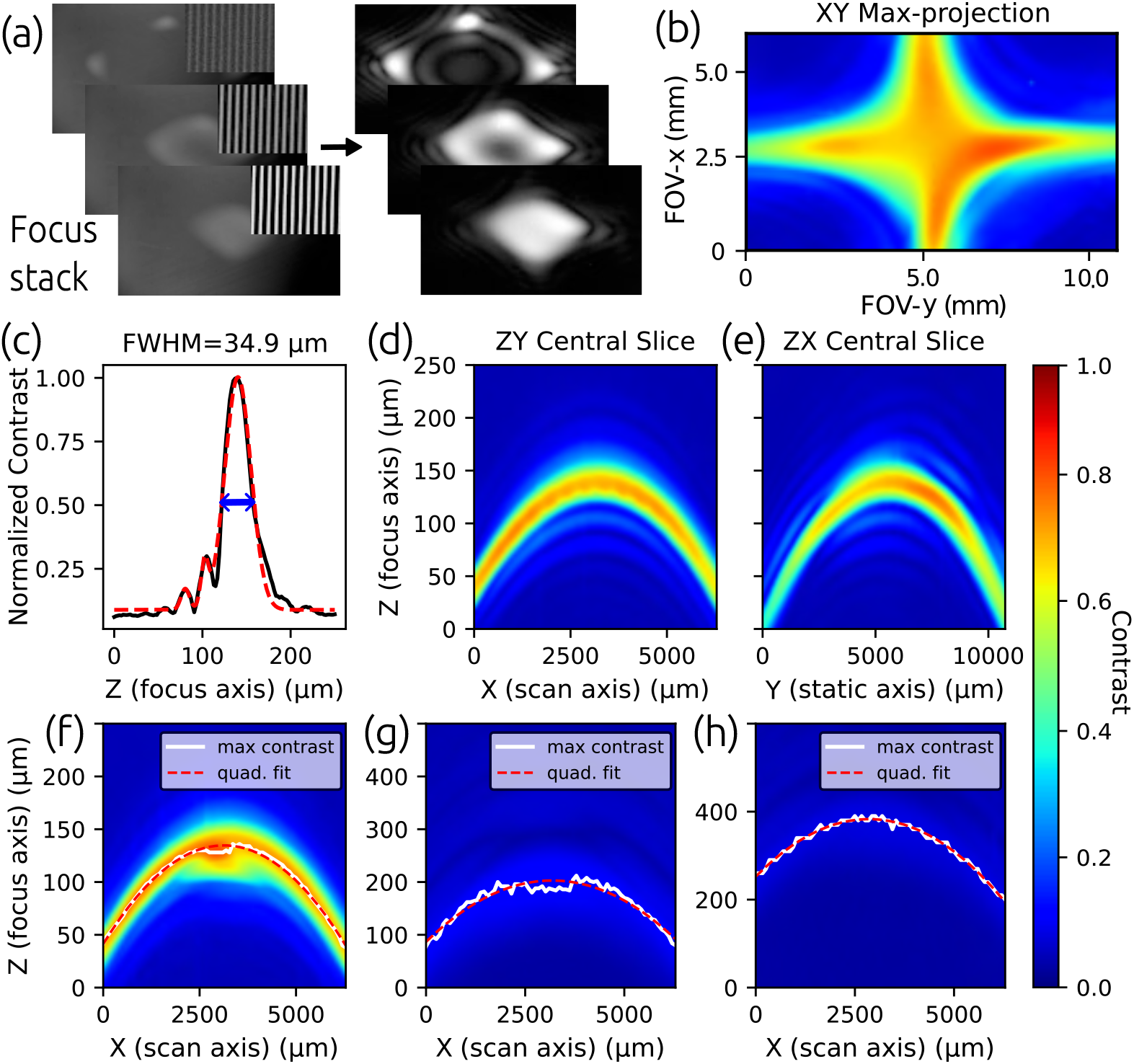
Field curvature characterization. (a) Three images from a representative z-stack of the Ronchi ruling using the full camera chip to characterize the change in contrast. (b) Contrast maximum projection along the focus axis measured in air for the entire camera chip. (c) In air depth-of-field calculation using the FWHM of the main lobe. (d) X-axis and (e) y-axis direction contrast slices along the light-sheet propagation axis. (f) Air, (g) water, (h) and oil maximum projections in the light-sheet volume (4 mm) along the y-axis. All heatmaps are normalized to the maximum contrast in air to illustrate the loss in contrast when imaging in different media. The maximum contrast and quadratic fits are shown in white and red respectively.

As the media’s refractive index increased, we observed increased spherical aberration and reduced contrast. For all images acquired in each media, we fit the central contrast slice along the focus axis with a multi-Gaussian function and calculated the effective depth of field (DOF) using the main lobe’s full width at half max (FWHM). Additionally, we visualized the expected contrast throughout the light-sheet volume by taking a max-projection over 4 mm around the center (Fig. 3, f-h). From these results, we estimated the region of maximum contrast follows a quadratic curvature with an overall axial shift of Δz ∼100 µm.

### 3.2. Quantifying increase in SBP using fluorescent microspheres

We validated our approach by imaging 1.0 µm fluorescent microspheres. While 1 µm microspheres are larger than the detection pathway NA, they still satisfy the effective resolution of the prototype curvedASLM when accounting for magnification and camera pixel size. We cropped the FOV to a height of 2125 µm by 6290 µm along the scan axis. We acquired 300 µm z-stacks with 0.5 µm axial steps. Microspheres were localized and fit using a 3D Gaussian PSF model to quantify image quality throughout the FOV [35]. curvedASLM increased the effective imaging range from 2500 µm to 6290 µm (Fig. 4). curvedASLM eliminated defocus aberrations; however, at the edges of the FOV, astigmatism remained, resulting in an asymmetric lateral stretching of the PSF.

**Fig. 4.**
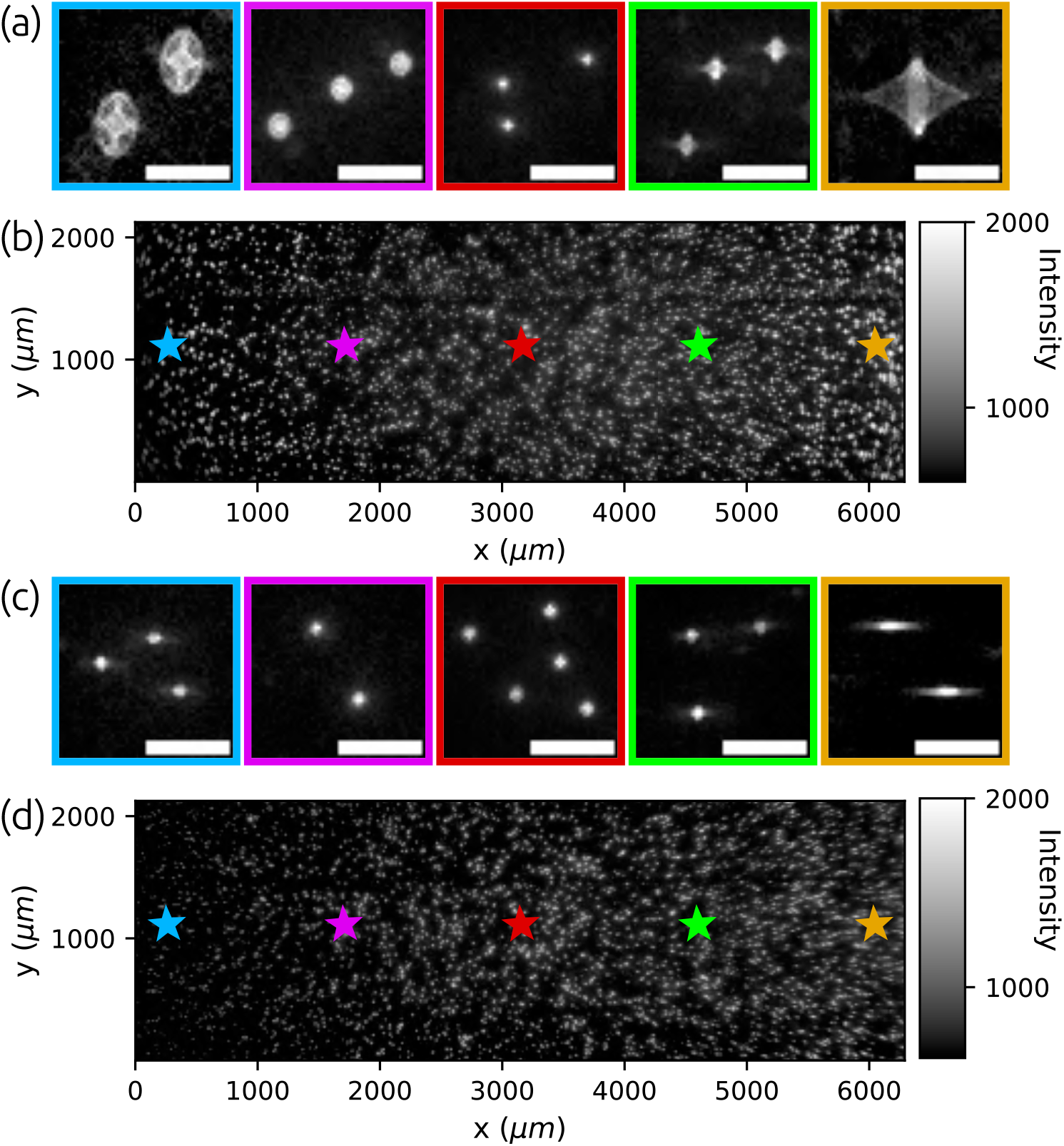
Usable FOV improvement with curvedASLM. 1.0 µm YG microspheres imaged at excitation wavelength *λ* =473 nm using the maximum FOV along the light-sheet propagation axis (2960 pixels or 6290 µm). (a,b) Conventional ASLM and (c,d) curvedASLM ROIs and full z-plane slices respectively. (a) Conventional ASLM and (c) curvedASLM ROIs highlighting the lateral PSF profile at throughout the FOV, the box colors correspond to the colored stars within (b,d). All scale bars are 50 µm.

Fig 5 summarizes the PSF fit results. The mean PSF size across the full camera FOV using curvedASLM was 2.8 µm x 2.5 µm (*σ*_xy_ × *σ*_z_), demonstrating that the curvature correction preserves subcellular resolution, while extending the usable imaging area by more than a factor of two. Due to residual aberrations, the range of spatially invariant image quality spans ∼5 mm, rather than the full camera width of 6290 µm.

**Fig. 5.**
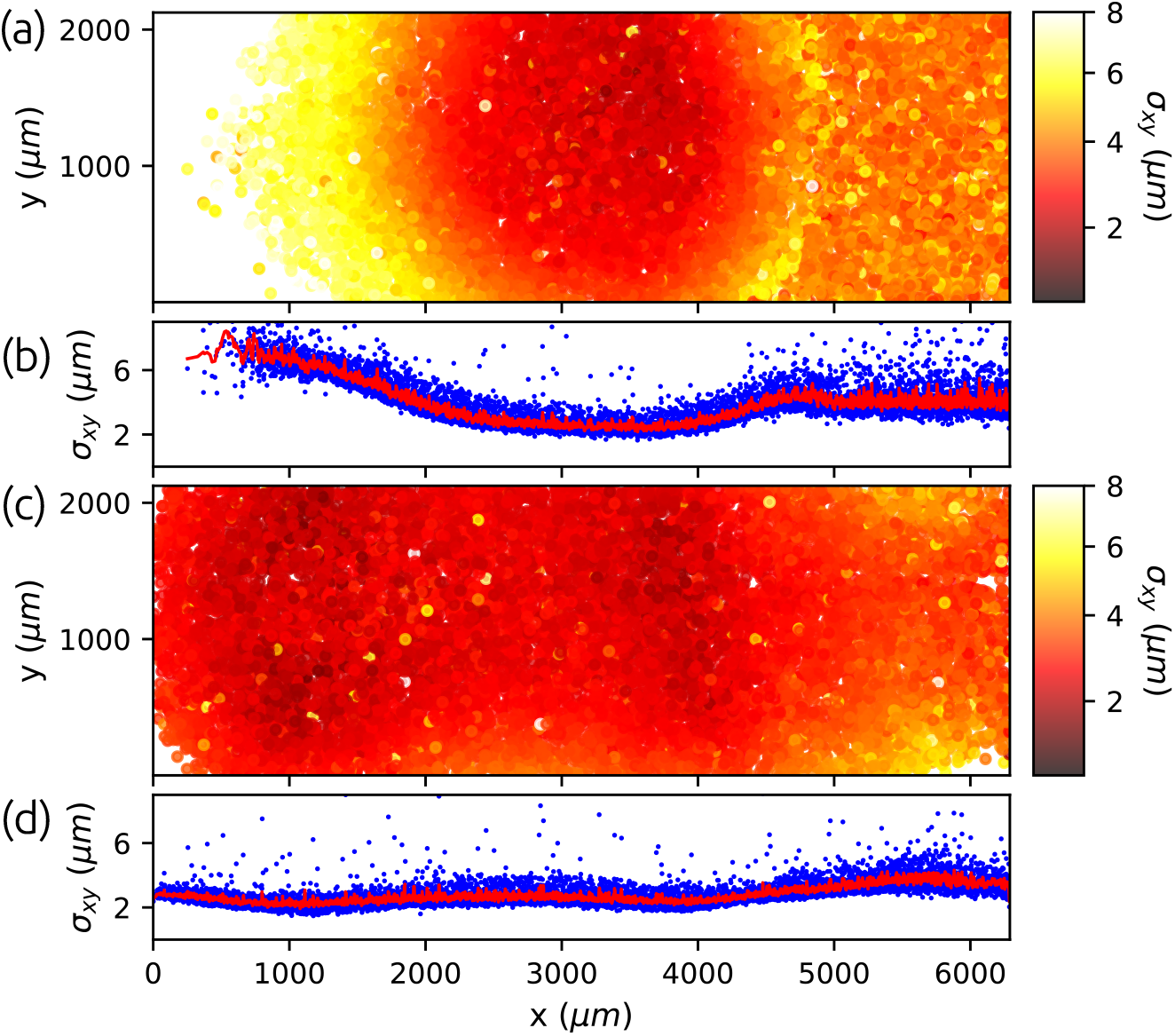
Lateral PSF improvement with curvedASLM. (a) Conventional ASLM and (c) curvedASLM heatmaps of *σ*_xy_ fit results for data in Fig. 4, illustrating the spatially varying image quality. (b) Conventional ASLM and (d) curvedASLM scatter plots of individual *σ*_xy_ (blue) fit results and rolling average (red).

### 3.3. Mouse brain imaging results

We evaluated the applicability of curvedASLM to large, heterogeneous biological samples by imaging the vascular network of a cleared mouse brain (Fig. 6). Using the full 6.29 mm field of view along the light-sheet propagation axis, we acquired the whole brain in six tiles.

**Fig. 6.**
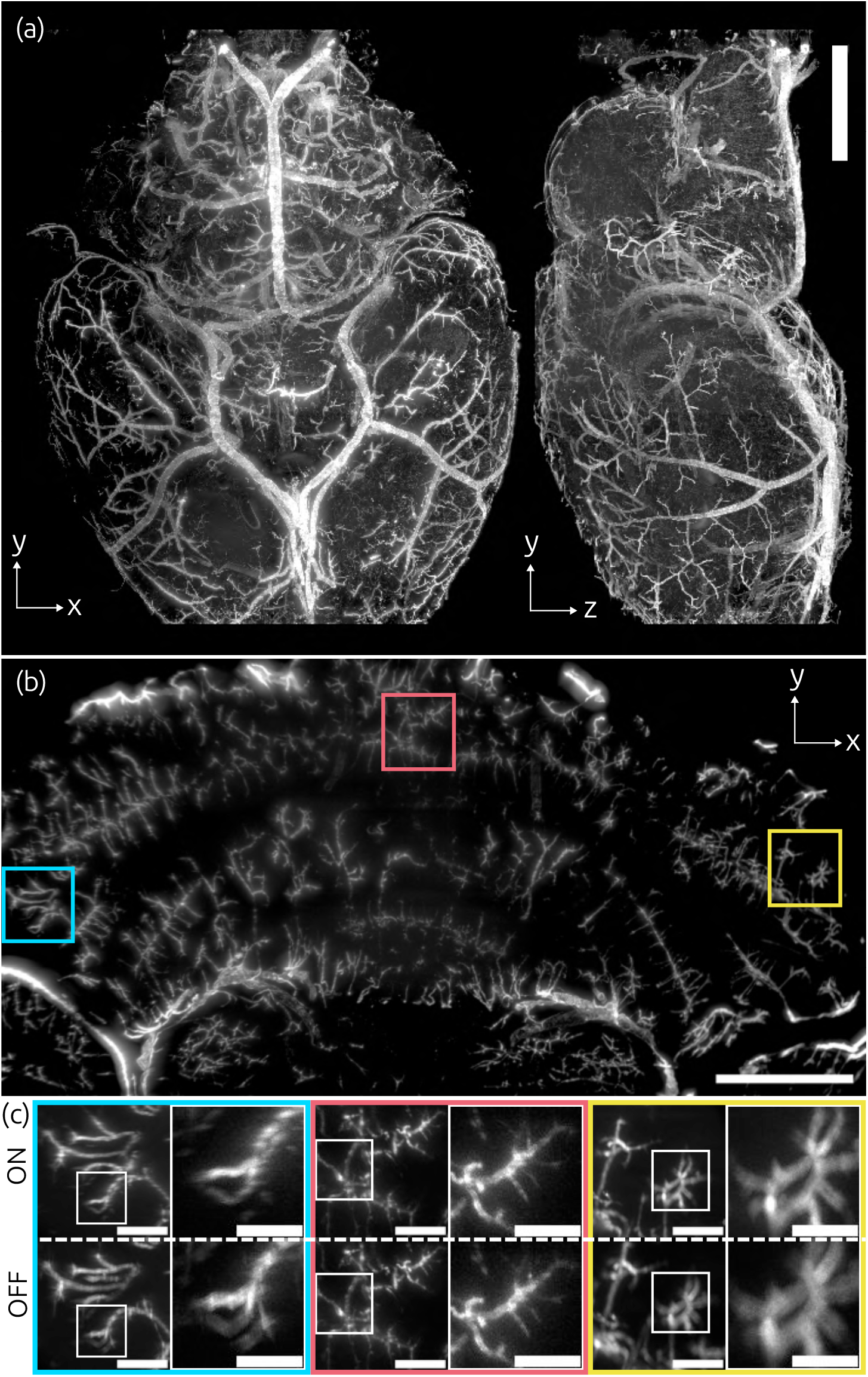
Cleared mouse brain vascular system imaging. (a) *XY* and YZ maximum projections, scale bar is 2.0 mm. (b) 400 µm *XY* projection sample data with ROI locations shown in (c), scale bar is 1 mm. (c) Cropped ROIs (531 µm ×531 µm) with (**ON**) and without (**OFF**) curvedASLM and enlarged subregions (200 µm ×200 µm). Scale bars in (c) are 200 µm and 100 µm for the enlarged ROIs.

To assess resolution uniformity across the extended FOV, we maximum projected a 3.19 mm×6.29 mm subregion over 400 µm, and sampled 531 µm square ROIs at the center and edges of the camera chip (Fig. 6b–c). Conventional ASLM exhibited characteristic blurring and loss of high-frequency content at the field edges, consistent with the mismatch between excitation and detection planes. We compared the raw data acquired with and without light-sheet curvature to illustrate the improved image quality, Fig. 6.

Python-based processing pipeline performed pixel shifting, multi-tile fusion, and optional de-striping for the full dataset. These results highlight that curvedASLM enables high-SBP imaging of large, cleared tissues with minimal hardware modifications and without sacrificing resolution or contrast across the field.

## 4. Discussion

Achieving a high SBP requires simultaneously maximizing the FOV, maintaining resolution, and properly digitizing the resulting analog image. Approaches to increase the SBP include designing custom and often bulky optics [14, 36], acquiring multiple images and stitching in either spatial [37] or frequency domains [38–40], or using curved imaging surfaces [22, 24]. However, to preserve optical sectioning and quality in LSFM, the independent, orthogonal optical pathways must remain co-planar across the entire FOV. As a result, even if the detection path supports a large SBP, misalignment between illumination and detection planes reduces the effective information capacity (Table 1).

It is important to note that the theoretical SBP values listed in Table 1 reflect the geometric FOV that satisfies the stated NA, and do not represent the realizable diffraction-limited FOV. Low-magnification, large-FOV objectives are typically diffraction-limited only near the optical axis. For example, the Olympus MVPLAPO 1×objective used in mesoSPIM and the MVPLAPO 2 ×used here maintain their specified NA across the entire image circle. However, off-axis aberrations confine diffraction-limited performance to the central region of the FOV. As a result, the realizable SBP is substantially smaller than the theoretical SBP implied by the objective specifications. We estimated the realizable diffraction-limited FOVs of the MVPLAPO 1× and 2× objectives (4.5 mm and 2.25 mm, respectively) using microsphere data acquired in an epifluorescence configuration without curvature correction [27]. By applying curvedASLM to correct field curvature, we increased the usable FOV to 6.3 mm. However, residual aberrations remain because these commercial objectives are not well corrected across their full FOV, preventing the full realization of their SBP potential. The distinction between theoretical and effective SBP underscores the need for methods such as curvedASLM, which restore image quality and extend the usable FOV of existing objectives.

In this work, we adapted the commonly used motorized light-sheet positioning mirror to dynamically adjust the light-sheet focus position during the synchronized light-sheet focus sweep in an ASLM platform. By applying a cubic waveform optimized for image content, curvedASLM dynamically corrects for field curvature. Our flexible approach is adaptable to any detection objective with a relatively simple calibration routine.

We demonstrate that curvedASLM can match the field curvature of a commercially available high-SBP objective with 2 ×magnification, 0.5 numerical aperture, and 2 cm working distance when imaging in air, water, or immersion oil. Previous attempts at using this objective for high-SBP LSFM and ASLM paired it with a zoom body with internal optics with apertures smaller than the large exit pupil (D_bfp_ ∼2 f NA =50 mm). As such, these approaches did not achieve the full performance of the objective. Instead of using the zoom body or a standard microscope tube lens, we instead used a 1× magnification and NA 0.25 objective (D_bfp_ ∼50 mm) to capture all of the light exiting the 2 ×objective and form an image on the camera sensor.

We evaluated system performance using fluorescent microspheres embedded in low-melt agar. Without curvature correction, the point spread function (PSF) away from the FOV center deviates significantly from the ideal shape due to defocus, leading to corrupted or missing data. We eliminated this dominant defocus aberration caused by the field curvature of the detection objective by applying light-sheet curvature during the ASLM sweep. After defocus removal, we observed residual aberrations, particularly astigmatism, at the edges of the FOV. Our findings match previously reported findings in a wide-field fluorescence microscope built using the same pair of high SBP objectives [27]. Recent work on spatially variant deconvolution may further improve the performance of the curvedASLM platform by removing residual aberrations [41, 42]. However, we chose not to implement deconvolution in this work because our images were not Nyquist-Shannon sampled due to the 4.25 µm pixel-pitch camera sensor. While these aberrations ultimately limit achievable image quality, there is still an associated brightness increase of nearly 25-fold when using the high-SBP objective here, as compared to a higher-magnification objective with similar NA (brightness B∝ NA^2^ / Mag^2^) [43].

The curvedASLM design presented here accounts only for field curvature along the light-sheet propagation direction. Improving the light-sheet generation pathway will further increase the achievable SBP by generating a taller light-sheet and utilizing more of the large-format camera chips [12, 14]. However, increasing the light-sheet size in the non-propagation direction will lead to the same field curvature issue, which can be mitigated by adapting the curvedLSFM light-sheet generation strategy to match the field curvature along non-propagating axis of the detection system [26]. Finally, adopting a large-format camera with smaller pixels (∼1.0 µm) that match the Nyquist-Shannon sampling criterion for NA_det_ = 0.5 and a programmable rolling shutter will increase the achievable resolution of the system and enable deconvolution.

Using a simple modification to the standard ASLM excitation pathway, curvedASLM solves the field curvature problem along one axis by dynamically optimizing the illumination volume to match the detection volume. We note that recent work demonstrated a static-field curvature correction for a high, isotropic-resolution ASLM [17] by mounting a curved mirror to the voice-coil unit used to translate the light-sheet focus. For high-speed applications, such a dedicated unit is essential due to the relatively slow response of electrotunable lenses. The trade-off for high-speed performance with a fixed mirror is flexibility in the choice of LSFM objective pair, whereas the curvedASLM approach presented here can correct field curvature for any LSFM objective pairing. An interesting future work would be to combine the two methods using transmissive adaptive optics [44] or other high-speed beam-steering and focusing strategies [45, 46].

Integrating the curvedASLM approach with the high-SBP detection objective used in this work increased the achievable FOV along the light-sheet propagation axis from 2500 µm to 6290 µm, representing a 2 ×increase in SBP. With the current sCMOS camera, the system is capable of 32-320 million voxels per second at typical exposure times of 10-100 ms. While this is lower than the ExA-SPIM and curvedLSFM voxel throughputs [14, 26], curvedLSFM has the potential to reach or surpass these voxel rates by replacing the camera with a model similar to the one used in the ExA-SPIM.

## Acknowledgments

The authors thank Dr. Adam Glaser (Allen Institute for Neural Dynamics) for guidance on high space-bandwidth product objective design and characterization.

## Data availability

The instrument control code is available at Ref. [29]. Ronchi ruler, fluorescent microsphere, and mouse brain ROI data are available at Ref. [47]. Image processing and analysis code is available at Ref. [48].

## References

1. H. Voie, D. Burns, and F. Spelman, “Orthogonal-plane fluorescence optical sectioning: Three-dimensional imaging of macroscopic biological specimens,” J. microscopy 170, 229–236 (1993).

2. J. Huisken, J. Swoger, F. Del Bene, et al., “Optical sectioning deep inside live embryos by selective plane illumination microscopy,” Science 305, 1007–1009 (2004).

3. R. M. Power and J. Huisken, “A guide to light-sheet fluorescence microscopy for multiscale imaging,” Nat. methods 14, 360–373 (2017).

4. M. B. Ahrens, M. B. Orger, D. N. Robson, et al., “Whole-brain functional imaging at cellular resolution using light-sheet microscopy,” Nat. methods 10, 413–420 (2013).

5. K. McDole, L. Guignard, F. Amat, et al., “In toto imaging and reconstruction of post-implantation mouse development at the single-cell level,” Cell 175, 859–876 (2018).

6. Kathe, M. A. Skinnider, T. H. Hutson, et al., “The neurons that restore walking after paralysis,” Nature 611, 540–547 (2022).

7. E. Sapoznik, B.-J. Chang, J. Huh, et al., “A versatile oblique plane microscope for large-scale and high-resolution imaging of subcellular dynamics,” Elife 9, e57681 (2020).

8. Mcfadden, Z. Marin, B. Chen, et al., “Adaptive optics in an oblique plane microscope,” Biomed. Opt. Express 15, 4498 (2024).

9. J. Park, D. J. Brady, G. Zheng, et al., “Review of bio-optical imaging systems with a high space-bandwidth product,” Adv. Photonics 3, 044001–044001 (2021).

10. K. M. Dean, P. Roudot, E. S. Welf, et al., “Deconvolution-free subcellular imaging with axially swept light sheet microscopy,” Biophys. journal 108, 2807–2815 (2015).

11. F. F. Voigt, D. Kirschenbaum, E. Platonova, et al., “The mesospim initiative: open-source light-sheet microscopes for imaging cleared tissue,” Nat. methods 16, 1105–1108 (2019).

12. N. Vladimirov, F. F. Voigt, T. Naert, et al., “Benchtop mesospim: a next-generation open-source light-sheet microscope for cleared samples,” Nat. Commun. 15, 2679 (2024).

13. Battistella, J. Schniete, K. Wesencraft, et al., “Light-sheet mesoscopy with the mesolens provides fast sub-cellular resolution imaging throughout large tissue volumes,” Iscience 25 (2022).

14. A. Glaser, J. Chandrashekar, S. Vasquez, et al., “Expansion-assisted selective plane illumination microscopy for nanoscale imaging of centimeter-scale tissues,” Elife 12, RP91979 (2025).

15. K. M. Dean, T. Chakraborty, S. Daetwyler, et al., “Isotropic imaging across spatial scales with axially swept light-sheet microscopy,” Nat. protocols 17, 2025–2053 (2022).

16. P. Ryan, E. A. Gould, G. J. Seedorf, et al., “Automatic and adaptive heterogeneous refractive index compensation for light-sheet microscopy,” Nat. communications 8, 612 (2017).

17. M. Aakhte, G. F. Müller, L. Roos, et al., “Isotropic, aberration-corrected light sheet microscopy for rapid high-resolution imaging of cleared tissue,” Nat. biotechnology pp. 1–11 (2025).

18. L. A. Barner, A. K. Glaser, L. D. True, et al., “Solid immersion meniscus lens (simlens) for open-top light-sheet microscopy,” Opt. letters 44, 4451–4454 (2019).

19. A. K. Glaser, N. P. Reder, Y. Chen, et al., “Multi-immersion open-top light-sheet microscope for high-throughput imaging of cleared tissues,” Nat. communications 10, 2781 (2019).

20. W. Feng, F. Zhao, F. Zhong, et al., “Flexible solid immersion meniscus lens (simlens) approach for enhancing biological imaging of cleared samples,” Opt. Lett. 49, 4126–4129 (2024).

21. S. J. Sheppard, P. T. Brown, and D. P. Shepherd, “Model based optimization for refractive index mismatched light sheet imaging,” Opt. Express 32, 36563–36576 (2024).

22. J. Fan, J. Suo, J. Wu, et al., “Video-rate imaging of biological dynamics at centimetre scale and micrometre resolution,” Nat. Photonics 13, 809–816 (2019).

23. M. Harfouche, K. Kim, K. C. Zhou, et al., “Imaging across multiple spatial scales with the multi-camera array microscope,” Optica 10, 471–480 (2023).

24. X. Yang, H. Chen, L. Kreiss, et al., “Curvature-adaptive gigapixel microscopy at submicron resolution and centimeter scale,” Opt. Lett. 50, 5977–5980 (2025).

25. Battistella, J. Schniete, K. Wesencraft, et al., “Light-sheet mesoscopy with the mesolens provides fast sub-cellular resolution imaging throughout large tissue volumes,” iScience 25, 104797 (2022).

26. L. Tang, J. Wang, J. Ding, et al., “Curved light sheet microscopy for centimetre-scale cleared tissue imaging,” Nat. Photonics pp. 1–8 (2025).

27. C. A. Werley, M.-P. Chien, and A. E. Cohen, “Ultrawidefield microscope for high-speed fluorescence imaging and targeted optogenetic stimulation,” Biomed. optics express 8, 5794–5813 (2017).

28. Z. Marin, X. Wang, D. W. Collison, et al., “Navigate: an open-source platform for smart light-sheet microscopy,” Nat. methods 21, 1967–1969 (2024).

29. Z. Marin, X. Wang, D. W. Collison, et al., “Navigate: Light-sheet control and acquisition software,” https://github.com/QI2lab/navigate/tree/develop (2025). Accessed: 2025-11-25; commit: 70ccd5a.

30. J. Huisken and D. Y. Stainier, “Even fluorescence excitation by multidirectional selective plane illumination microscopy (mspim),” Opt. letters 32, 2608–2610 (2007).

31. J. Moore, D. Basurto-Lozada, S. Besson, et al., “Ome-zarr: a cloud-optimized bioimaging file format with international community support,” Histochem. Cell Biol. 160, 223–251 (2023).

32. M. Albert, A. Wilhelmi, J. de Folter, et al., “multiview-stitcher,” (2025). https://github.com/multiview-stitcher/multiview-stitcher.

33. T. Peng, Y. Liu, G. Müller, et al., “Leonardo: A toolset to remove sample-induced aberrations in light sheet microscopy images,” bioRxiv (2025).

34. N. Renier, Z. Wu, D. J. Simon, et al., “iDISCO: a simple, rapid method to immunolabel large tissue samples for volume imaging,” Cell 159, 896–910 (2014).

35. P. T. Brown, S. Sheppard, A. Coullomb, and D. P. Shepherd, “localize-psf,” (2023). https://github.com/QI2lab/localize-psf.

36. McConnell J. Trägårdh, R. Amor, et al., “A novel optical microscope for imaging large embryos and tissue volumes with sub-cellular resolution throughout,” Elife 5, e18659 (2016).

37. D. Hörl, F. Rojas Rusak, F. Preusser, et al., “Bigstitcher: reconstructing high-resolution image datasets of cleared and expanded samples,” Nat. methods 16, 870–874 (2019).

38. Zheng, R. Horstmeyer, and C. Yang, “Wide-field, high-resolution fourier ptychographic microscopy,” Nat. photonics 7, 739–745 (2013).

39. M. G. Gustafsson, “Surpassing the lateral resolution limit by a factor of two using structured illumination microscopy,” J. microscopy 198, 82–87 (2000).

40. X. Ou, R. Horstmeyer, G. Zheng, and C. Yang, “High numerical aperture fourier ptychography: principle, implementation and characterization,” Opt. express 23, 3472–3491 (2015).

41. K. Yanny, K. Monakhova, R. W. Shuai, and L. Waller, “Deep learning for fast spatially varying deconvolution,” Optica 9, 96–99 (2022).

42. A. Kohli, A. N. Angelopoulos, D. McAllister, et al., “Ring deconvolution microscopy: exploiting symmetry for efficient spatially varying aberration correction,” Nat. Methods pp. 1–10 (2025).

43. M. Born and E. Wolf, Principles of optics: electromagnetic theory of propagation, interference and diffraction of light (Pergamon Press, 1975). Section 9.8.3.

44. P. Pozzi, M. Quintavalla, A. Wong, et al., “Plug-and-play adaptive optics for commercial laser scanning fluorescence microscopes based on an adaptive lens,” Opt. Lett. 45, 3585–3588 (2020).

45. M. Duocastella, G. Sancataldo, P. Saggau, et al., “Fast inertia-free volumetric light-sheet microscope,” ACS Photonics 4, 1797–1804 (2017).

46. M. R. Rai, C. Li, and A. Greenbaum, “Quantitative analysis of illumination and detection corrections in adaptive light sheet fluorescence microscopy,” Biomed. Opt. Express 13, 2960–2974 (2022).

47. S. J. Sheppard, “curvedaslm datasets,” (2024). 10.5281/zenodo.17717241.

48. D. S. P. Sheppard, Steven J., “curvedaslm support code,” (2025). 10.5281/zenodo.17717435.

